# Exact *p*-values for global network alignments via combinatorial analysis of shared GO terms

**DOI:** 10.1101/2020.10.08.332254

**Authors:** Wayne B. Hayes

## Abstract

Network alignment aims to uncover topologically similar regions in the protein-protein interaction (PPI) networks of two or more species under the assumption that topologically similar regions perform similar functions. Although there exist a plethora of both network alignment algorithms and measures of topological similarity, currently no “gold standard” exists for evaluating how well either is able to uncover functionally similar regions. Here we propose a formal, mathematically and statistically rigorous method for evaluating the statistical significance of shared GO terms in a global, 1-to-1 alignment between two PPI networks. We use combinatorics to precisely count the number of possible network alignments in which *k* proteins share a particular GO term. When divided by the number of all possible network alignments, this provides an explicit, exact *p*-value for a network alignment with respect to a particular GO term.

## 1 Introduction

Network alignment aims to uncover similar network connection patterns between two or more networks under the assumption that common network topology (which may be easily observable) correlates with common function (which is more difficult to observe).

In this paper we focus on the simplest case to analyze: a pairwise global network alignment (PGNA). Pairwise, because exactly two networks are being aligned; and global, because we insist that every node in the smaller network (ie., the one with fewer nodes) is mapped to exactly one node in the larger network. Extending the analysis to the non-1-to-1 case—as well as to local and multiple network alignments—appears nontrivial and is left for future work.

Network alignment algorithms abound and their number is increasing rapidly. There is an urgent need for a *rigorous* way to evaluate these methods for their ability to identify functionally similar proteins across species. Unfortunately, there are almost as many ways of evaluating network alignments as there are alignment algorithms. The research area is severely hampered by the lack of any “gold standard” for evaluating the functional similarity that is uncovered by any given network alignment. One of the most common methods for evaluating biological significance involves using the Gene Ontology’s GO term hierarchy. While other biologically relevant information exists, such as KEGG pathways (any others?), in this paper we focus on leveraging GO terms to produce a mathematically and statistically rigorous “gold standard” measure of the statistical significance of an alignment in which exactly *k* aligned protein pairs all share a common GO term *g*.

## 2 Method: GO-term *p*-values by exhaustive enumeration of alignments

### 2.1 Network alignment and functional similarity

Given two networks *G*_1_, *G*_2_, let the node sets *V*_1_, *V*_2_ represent *n*_1_ and *n*_2_ proteins respectively, and the edge sets *E*_1_, *E*_2_ represent protein-protein interactions (PPIs). Assuming (without loss of generality) that *n*_1_ ≤ *n*_2_, a pairwise global network alignment (PGNA) is a 1-to-1 mapping *f*: *V*_1_ → *V*_2_ in which every node in *V*_1_ is mapped to exactly one node in *V*_2_.

Once an alignment is specified, we need to measure the extent to which functionally similar proteins have been mapped to each other. Many existing methods evaluate their alignments using GO terms from the Gene Ontology [1], and most often evaluate the functional similarity of each pair of aligned proteins independent of all the others, and then average across the pairs. While the score of each pair may be meaningful, taking an average assumes that each pair is independent of all the others. This is not true because the pairings themselves are inter-dependent via the alignment itself, which is built globally. For example, in a 1-to-1 alignment, each node in each network can appear at most once across the entire alignment, a property which destroys the independence assumption needed for a meaningful average across aligned node pairs.

There are other considerations as well: most GO terms have inter-dependencies with most other GO terms via the GO hierarchy [2]; most proteins have more than one GO annotation, and it is difficult to create a measure that correctly evaluates similarity between two proteins with different sets of GO terms that only partially overlap; and finally, since most GO terms annotate many proteins, it is nontrivial to compute the significance of aligning a set of protein pairs while accounting for both the frequency and inter-relationships between GO terms that may appear in multiple pairs across the set of aligned pairs.

Existing solutions to evaluate the relationships between GO terms, as well as between the gene products that they annotate, are currently all *ad hoc*[2]. Specifically in the context of network alignment, to our knowledge all of the existing methods are also *ad hoc*, and lack any strong mathematical or statistical rigor. (The details of these other methods are irrelevant and entirely unrelated to our work, and so are relegated to the Appendix, though we offer a discussion of their pitfalls in Section §4.) If we are to make progress in the network alignment community, we need a way to compare network alignments for the extent to which they uncover functional similarity among its aligned proteins. Given the plethora of *ad hoc* methods currently being used, there is an urgent need for a “gold standard” measure of the significance of functional similarity uncovered by a network alignment. Here we propose such a gold standard, which is both mathematically and statistically rigorous.

### 2.2 Computing the total number of possible alignments

In the following exposition, we must discuss in great detail the combinatoric structure of a given alignment. To aid visualization, we use what I call the “Pegs and Holes” analogy: given networks *G*_1_, *G*_2_ with *n*_1_, *n*_2_ nodes, we imagine *G*_2_’s nodes as *n*_2_ identical “holes” drilled into a large board, and *G*_1_’s nodes as *n*_1_ identical “pegs” that can each fit into any hole. To enforce the global 1-to-1 property, there are two cases:

1. *n*_1_ ≤ *n*_2_, so every peg is placed into some hole, leaving *n*_2_ – *n*_1_ empty holes. There are 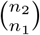 ways to choose which holes to use, and *n*_1_! ways to place the pegs.
2. *n*_1_ > *n*_2_, so every hole is filled with some peg, leaving *n*_1_ – *n*_2_ pegs unplaced. There are 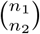 ways to choose which pegs to place, and *n*_2_! ways to place them.

The above two cases are symmetric and so, without loss of generality, we assume *n*_1_ ≤ *n*_2_. Then, the total number of all possible alignments is

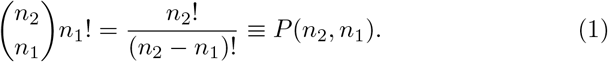

The function *P*(·, ·) of Eq. (1) is more commonly known as *k-permutations-of-n*, or *P*(*n, k*). However, *P*(*n, k*) is usually defined to be zero if *n* < *k*, whereas we will often need to compute the number of alignments when we don’t know which of the two values is larger. Thus, in this paper, we will adopt a modified permutation function *π*(*n*_1_, *n*_2_) as follows

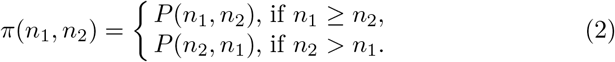

### 2.3 Counting alignments with exactly *k* matches

Given a particular GO term *g*, assume *g* annotates λ_1_ pegs and λ_2_ holes. A peg and the hole it sits in are, more technically, a pair of aligned nodes. We say that such a pair “match” with respect to GO term *g* if they are both annotated with *g*. Let 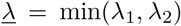, and 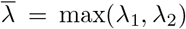. Given a random 1-to-1 alignment, we are going to compute the probability *p* that exactly *k* pairs of aligned nodes share *g*. In our analogy, this means that exactly *k* pegs—no more, no less—that are annotated with *g* sit in holes that are also annotated with *g*. To do this, we will use a combinatorial argument to enumerate all possible PGNAs that can exist that have exactly *k* matches. Given that number, we simply divide by Eq. (1) to get the probability that a randomly chosen alignment has exactly *k* matches.

#### 2.3.1 Special cases

The following are special cases:

1. if 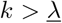, then *p* = 0.
2. if 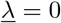, then *p* = 1 if *k* = 0 and *p* = 0 otherwise.
3. if λ_2_ = *n*_2_, then *p* = 1 if *k* = λ_1_, and *p* = 0 otherwise.
4. if λ_1_ > *n*_2_ – λ_2_ and *k* < λ_1_ – (*n*_2_ – λ_2_), then *p* = 0, otherwise *p* > 0 is computed below.

The last case arises when λ_1_ > *n*_2_ – λ_2_, which means that there are more annotated pegs than non-annotated holes, necessitating that *at least* λ_1_ – (*n*_2_ – λ_2_) annotated pegs must align with annotated holes. (Recall we are computing the probability of *exactly k* aligned pairs sharing *g*, so *k* too small in this case gives *p* = 0.)

Below we describe the general case in detail. In broad outline, there are three steps: (i) create the required *k* matches by placing *k* annotated pegs into *k* annotated holes; (ii) arrange to place the remaining annotated pegs away from the annotated holes in order to keep *k* constant; (iii) place any remaining pegs (all of which are non-annotated) in any still-empty holes (some of which may be annotated). In each case we either sum, or multiply, as appropriate, the number of ways to perform the described action. In the end we have counted all the possible ways to create an alignment that has exactly *k* matches.

#### 2.3.2 Creating exactly k matches

Out of the λ_1_ pegs annotated with *g*, pick 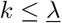 of them; there are 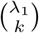 ways to do this. We will place these *k* pegs into *k* holes that are also annotated with *g*; there are 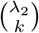 ways to pick the holes, and *k*! ways to place the *k* pegs into the *k* holes. Thus, the total number of ways to match exactly *k* pairs of nodes that share *g* is

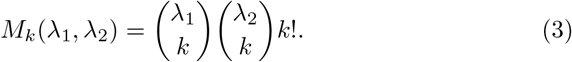

From this point onward, in order to keep k constant, we are committed to creating no more matches.

#### 2.3.3 Enumerating the ways to use the remaining annotated holes

To ensure that no more node pairs are matched, we need to ensure that none of the remaining (λ_1_ – *k*) annotated pegs are placed into any of the remaining (λ_2_ – *k*) annotated holes. Thus, each annotated hole must either remain empty, or take an non-annotated peg. There are *n*_1_ – λ_1_ available non-annotated pegs, regardless of the value of *k*. Pick *μ* of them. Since these *μ* pegs are all non-annotated, they can go into any unoccupied annotated hole without changing *k*. However, there are lower and upper bounds on what *μ* can be, as follows:

– *μ* can be at most 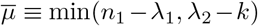, since *n*_1_ – λ_1_ is the total number of non-annotated pegs, and λ_2_ – *k* is the number of available annotated holes in which to place (some of) them.
– note that we have *n*_1_ – *k* pegs (of both types) remaining to place, and exactly *n*_2_ – λ_2_ non-annotated holes, into which some (or all) of the pegs can be placed. By the pigeon hole principle, if (*n*_1_ – *k*) > (*n*_2_ – λ_2_), then some of the pegs—and they can only be non-annotated pegs—*must* go into annotated holes. Thus, *μ*—which refers only to non-annotated pegs—must be at least 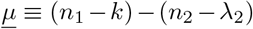 if (*n*_1_ – *k*) > (*n*_2_ – λ_2_); otherwise 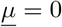.

#### 2.3.4 Distributing the remaining pegs

For any 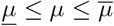, we need to count how many alignments can be built when *μ* non-annotated pegs are placed into the λ_2_ – *k* available annotated holes, as well as what happens to all the remaining pegs. The process is as follows.

1. There are 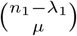 ways to choose *μ* non-annotated pegs, and *π*(λ_2_ – *k, μ*) ways to align them with the open annotated holes. To simplify notation note that *n*_1_, *n*_2_, λ_1_, λ_2_ are all fixed; thus, let 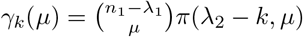.
2. Recall that there are still λ_1_ – *k* annotated pegs to be placed, and that they must be placed into non-annotated holes, so we must “reserve” λ_1_ – *k* non-annotated holes, which will be further accounted for below.
3. Once *μ* annotated holes are filled with non-annotated pegs, the rest of the annotated holes must remain empty; this leaves *n*_1_ – λ_1_ – *μ* non-annotated pegs to go into the *n*_2_ – λ_2_ non-annotated holes. Keeping in mind the “reservation” above, there are *n*_2_ – λ_2_ – (λ_1_ – *k*) available non-annotated holes. There are 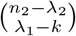 ways to choose which holes to use while reserving λ_1_ – *k* of them, and *π*(*n*_1_ – λ_1_ – *μ, n*_2_ – *λ*_2_ – (λ_1_ – *k*)) ways to place the pegs into the chosen holes; let 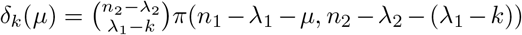.
4. Finally, we place the remaining λ_1_ – *k* annotated pegs into the reserved holes of the same number; there are (λ_1_ – *k*)! ways to do this.

#### 2.3.5 Summing the unmatched region of the alignment

Combining all of the above for fixed *μ* and then summing over all possible *μ*, the total number of ways that *n*_1_ – λ_1_ non-annotated pegs can be used to (partially or wholly) fill λ_2_ – *k* annotated holes, and then use all the remaining pegs and holes in a manner consistent with keeping *k* constant, is

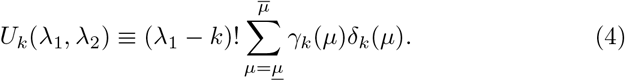

#### 2.3.6 Final tally for exactly k matches

Combining Eq.s (3) and (4), the total number of alignments in which exactly k aligned node pairs share GO term *g* is

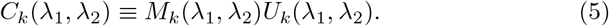

### 2.4 The probability of an alignment with exactly k matches

Eq. (5) counts all possible alignments in which exactly *k* aligned node pairs share GO term *g*. To get the probability *p_k_* of the same event, we divide by Eq. (1):

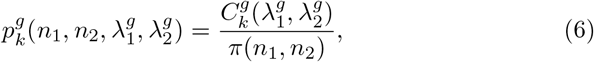

where a superscript *g* has been added as appropriate to denote that this probability is specifically tied to GO term *g*.

Note this refers to *exactly k* matches. To measure the statistical significance of *m* matches, we sum Eq. (6) for *k* from *m* to 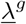.

### 2.5 Efficiently dealing with huge numbers

Though technically it is only an implementation detail, it is important to briefly discuss how to deal with the astronomically huge numbers involved in these calculations. Typical modern biological networks can have thousands to tens of thousands of nodes, and some GO terms annotate thousands of genes in each network. For example, in BioGRID 3.4.164 that we use below, the two biggest PPI networks in terms of number of nodes are *H. sapiens* and *A. thaliana*, which contain exactly 17,200 and 9,364 unique proteins, respectively, that are involved in physical interactions. Eq. (1) in this case is approximately 10^38270^—an integer with over 38,000 digits in base-10, which is far above the values typically representable on modern hardware. Luckily, its logarithm is easy to represent in double precision floating point, and so all of the multiplications herein can be computed as the floating-point sum of logarithms. The sole complication is the summation in Eq. (4), which is a sum of *values*, not logarithms. We use the following trick. Given two numbers *a* and *b*, assume we have at our disposal only their logarithms, *α* = log(*a*) and *β* = log(*b*). Our goal is to estimate log(*a* + *b*). Without loss of generality, assume *a* ≤ *b*. Then,

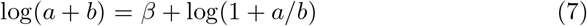

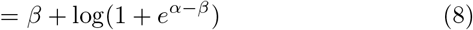

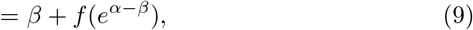

where *f*(*x*) is some function that can provide an accurate estimate of log(1+*x*) for any |*x*| ≤ 1. One must be careful because if |*x*| is below the machine epsilon (≈ 10^-16^ in double precision), then 1 + *x* evaluates to 1 because *x* is rounded away, and a direct evaluation of the expression log(1 + *x*) gives zero. The solution is not hard: the built-in library function for log can evaluate log(1 + *x*) with sufficient accuracy if |*x*| > 10^-6^; for smaller values of |*x*|, we explicitly invoke the Taylor series, which is extremely accurate for small values of |*x*|. We have tested that this method gives values for log(*a* + *b*) that are accurate to almost machine precision for any |*x*| ≤ 1.

## 3 Results

### 3.1 Numerical Validation

Staring at *C_k_*(λ_1_, λ_2_) in Eq. (5) and tracing back through the equations that define its components, it is not immediately obvious that the *C_k_*(λ_1_, λ_2_), when summed over all possible values of k, must add up to exactly *π*(*n*_1_, *n*_2_) independent of the choice of λ_1_, λ_2_. Yet if Eq. (5) is correct, then this must be the case since summing *p_k_* in Eq. (6) across all *k* of must give exactly 1.

In the calculation of 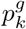 in Eq. (6), the values of *k* and *g* are fixed. For a fixed *g*, valid values of *k* range from zero to 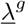. If our calculations are correct, then the sum across *k* of 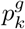 should be exactly 1 for any fixed *g, n*_1_, *n*_2_, λ_1_, λ_2_. We tested tested this property in the following cases:

1. exhaustively for all 0 ≤ λ_1_ ≤ *n*_1_ and 0 ≤ λ_2_ ≤ *n*_2_ for all 0 ≤ *n*_1_ ≤ *n*_2_ ≤ 100;
2. as above but in steps of 10 in λ_*i*_ and *n_i_* up to *n*_2_ = 1, 000;
3. as above but in powers of 2 in λ_*i*_ and *n_i_* up to *n*_2_ = 32, 768;
4. several billion random quadruples of (*n*_1_, *n*_2_, λ_1_, λ_2_) with *n*_2_ chosen uniformly at random up to 100,000, *n*_2_ chosen uniformly at random up to *n*_2_, and the λ’s chosen uniformly at random up to their *n* value.

We found in all cases that the difference from 1 of the sum over *k* of 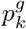 was bounded by 10^-9^. (Keep in mind that we had access only to the logarithms of the *C_k_*; that the actual sum across *k* had to be approximated term-by-term using Eq. (9); that the correct answer in log space is log(1) = 0; and that all operations were performing in floating point, which incurs roundoff error.) Furthermore, in any particular case, the numerical (floating-point roundoff) error will be dominated by the sum over *μ* in Eq. (4), and so we would expect the error to be smaller (ie., sum closer to 1) when there are fewer terms in Eq. (4). The number of terms is well-approximated by min(*n*_1_ – λ_1_, *n*_2_). Indeed, we find that if the sum was *S*, then the value |*S* – 1|/min(*n*_1_ – λ_1_, *n*_2_) has mean ≈ 3 × 10^-14^, standard deviation ≈ 3 × 10^-13^, and was never observed to exceed 3 × 10^-12^.

### 3.2 Validation against random alignments of real PPI networks

We downloaded the 8 largest protein-protein interaction networks from release 3.4.164 of BioGRID (cf. Table 1), and the GO database release of the same month. As many authors of network alignment papers do, we then split the GO database into two versions: one with all GO terms, and ones where sequence-based GO terms were disallowed. For each of the 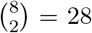 pairs of networks and for both versions of the GO database, we generated 400 million random alignments, for a total of 22.4 billion random alignments. For each GO term *g*, we observed the integer frequency 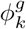 that *g* was shared by exactly *k* proteins when it annotated 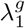 out of *n*_1_ proteins in network *G*_1_ and 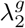 proteins out of *n*_2_ in network *G*_2_. (Note that formally 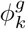 it has six parameters, 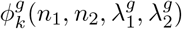), though we often abbreviate it to 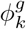 or even just *ϕ_k_* or *ϕ* if context is clear.) It is a non-negative integer bounded by the number of random alignments, *N* = 4 × 10^8^, and dividing it by *N* gives an estimate of the probability that a randomly chosen alignment between *G*_1_ and *G*_2_ will contain exactly *k* aligned protein pairs that share *g*.

**Table 1.** 8 largest networks of BioGRID 3.4.164, sorted by node count.

| nodes | common name | official name |
| --- | --- | --- |
| 17200 | Human | *H. sapiens* |
| 9364 | Thale cress | *A. thaliana* |
| 8728 | Fruit fly | *D. melanogaster* |
| 6777 | Mouse | *M. musculus* |
| 5984 | Baker's yeast | *S. cerevisiae* |
| 3194 | Worm | *C. elegans* |
| 2811 | Fission yeast | *S. pombe* |
| 2391 | Rat | *R. norvegicus* |

The estimated (ie., observed) probability 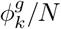 can be compared to 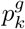 of Eq. (6). Across the 22.4 billion random alignments, we observed 428,849 unique combinations of the six parameters 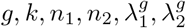 that formally define 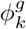. Figure 1 is a scatter plot of 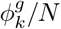 for all 428,849 of them, versus the theoretical value from Eq. (6). The agreement is excellent.

**Fig. 1.**
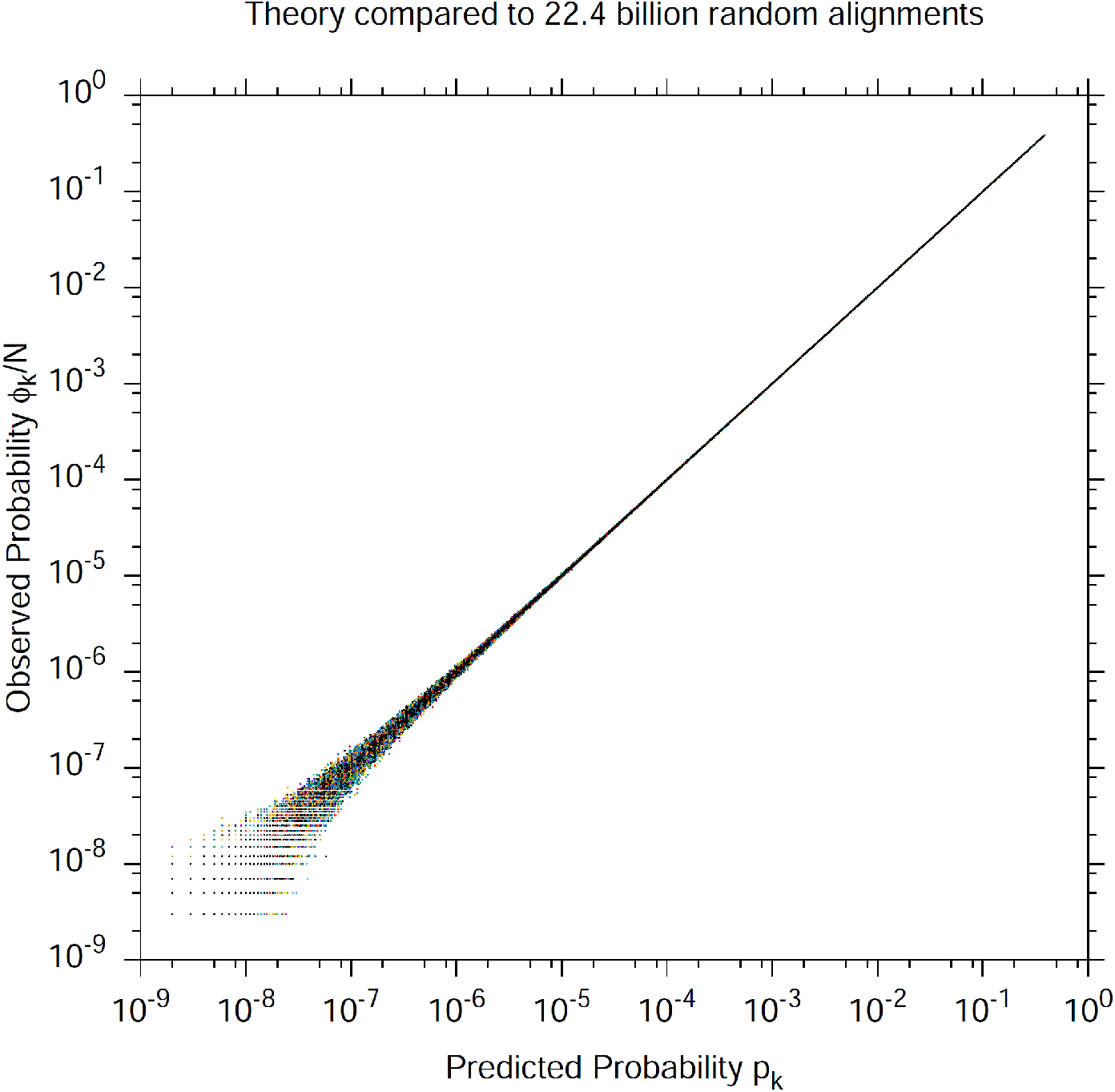
Scatter plot of the observed *ϕ_k_*/*N* vs. theoretical *p_k_* probability across 22.4 billion random alignments between pairs of networks from BioGRID 3.4.164. The vertical axis depicts the observed probability of an event, which is the observed frequency 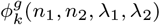 divided by the number of samples *N* = 4 × 10^8^. The horizontal axis is the value given by Eq. (6) for the parameters of the observation. There are 428,849 observations plotted across all observed values of 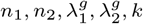.

The scatter in Figure 1 increases towards the low end because events with probability near *N*^-1^ are rarely observed, and so the estimate of their probability contains significant sampling noise. In fact there is “width” to the scatter plot at all values of probability, but it is difficult to observe in Figure 1. To more clearly see the scatter, we compute the *ratio* of the observed to theoretical values of probability, which will have an expected value of 1 if Eq. (6) is an accurate and unbiased estimator of probability. Figure 2 plots the mean and standard deviation (binned in powers of 2 of the number of samples) of 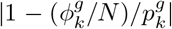 across all 428,849 observed frequencies, as a function of the number of samples that gave rise to the probability estimate. We can clearly see that the ratio approaches 1 asymptotically with the square root of the number of samples, consistent with sampling noise in *ϕ*.

**Fig. 2.**
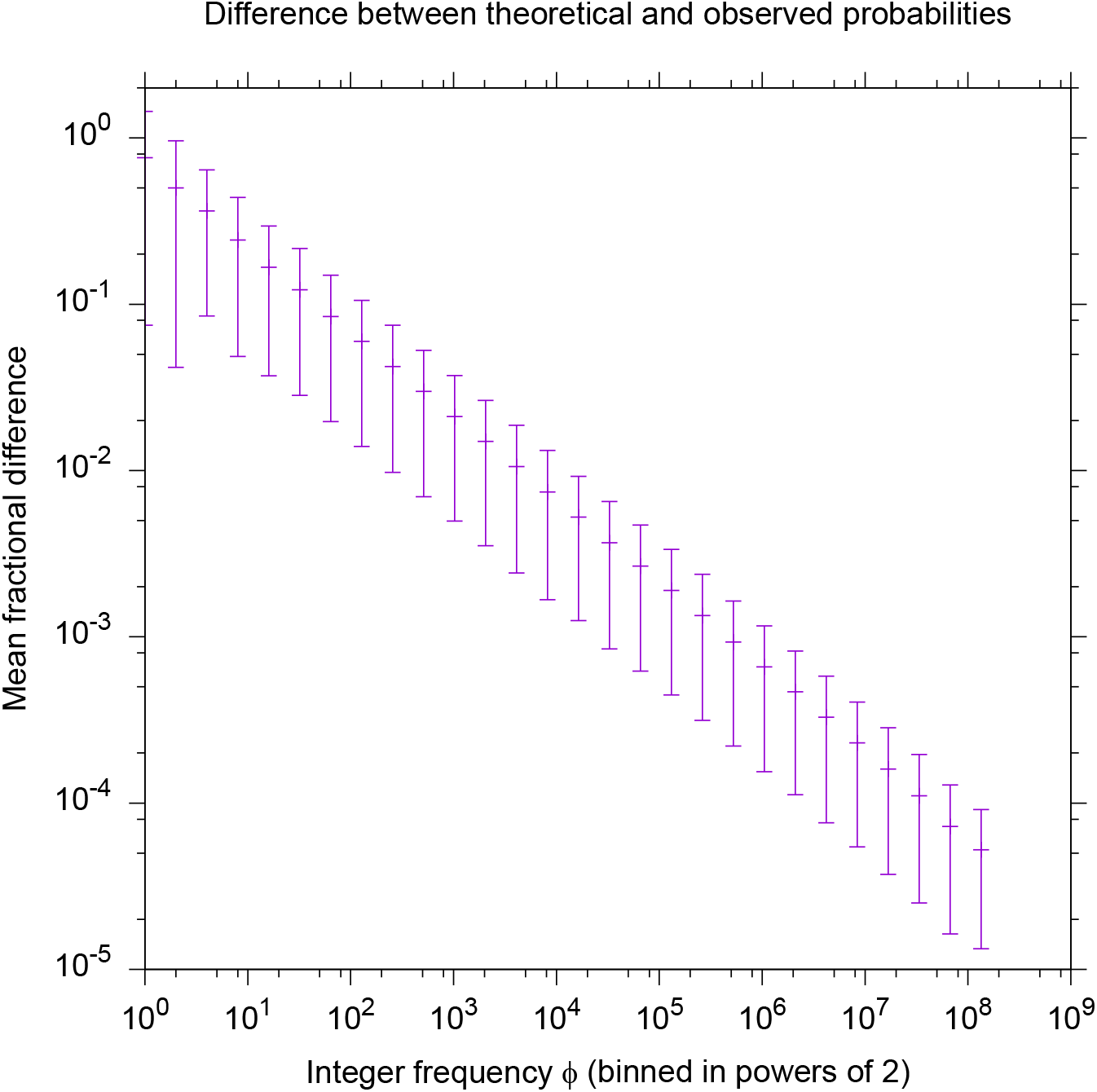
Same data as Figure 1, except that, for each point, we have computed the distance *D* from 1 of the ratio of observed to predicted probability: 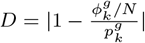. Each observed frequency 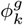 (which we will henceforth abbreviate a *ϕ*) is converted to an observed probability *ϕ/N*, where *N* is the number of random alignments (4 × 10^8^) per pair of networks. However, *ϕ* is also the number of samples used to create the observed probability estimate; higher *ϕ* gives a better estimate of the probability. We binned *ϕ* in powers of 2 (ie. the bin is ⌊log_2_(*ϕ*)⌋, and for each bin plotted the mean and standard deviation of *D*. We see that as the number of samples increases, the ratio approaches 1 as the square root of the number of samples, consistent with sampling noise.

### 3.3 Comparison with a simpler Poisson model

We introduce a Poisson-based model that correctly iterates across GO terms rather than protein pairs, though it simplistic and only provides an approximate *p*-value. Given that GO term *g* annotates λ_*i*_ out of *n_i_* proteins in network *G_i_*, then a randomly chosen protein *u_i_* from network *G_i_* has probability λ_*i*_/*n_i_* of being annotated by *g*. Thus, when a pair of proteins (*u*_1_, *u*_2_) is independently sampled from all possible pairs of proteins in *V*_1_ × *V*_2_, the probability that they both share *g* is 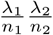; note that, at this stage, this *not* an approximation—the probability is exact. Since multiple independent Poisson processes have a cumulative rate which is simply the sum of their individual rates, if we choose *m* such pairs of nodes, *each independently of all the others*, then the number of pairs that share *g* is modeled by a Poisson distribution with mean 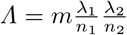, and the probability that *k* such pairs share *g* is

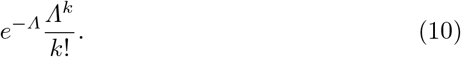

While this distribution correctly models the case where each protein pair is chosen independently and uniformly at random from all the others, it is only an approximation for the distribution of protein pairs that share *g* in a 1-to-1 network alignment, because the set of node pairs in an alignment are *not* independent: each pairs depends implicitly on all the others via the alignment itself, which is built globally and disallows any one node from being used more than once.

In relation to the GO term frequencies, Eq. (10) is a good approximation when λ_1_ ≪ *n*_1_ and λ_2_ ≪ *n*_2_, because then the probabilities are small and each pair that shares *g* has only a small influence on others. However, the approximation gets worse as either of λ_1_ or λ_2_ increases. To demonstrate this, we took an assortment of *good* alignments between the 3.4.164 BioGRID networks [3] which had some astronomically small *p*-values. Figure 3 plots the ratio of the Poisson-based *p*-value of Eq. (10) to the correct one of Eq. (6), as a function of λ_1_ + λ_2_. As we can see, the ratio is rarely less than 1 (ie., the Poisson-based *p*-value is almost always greater than or equal to the exact one), but can be huge if λ_1_ + λ_2_ is large—meaning the Poisson model grossly underestimates the statistical significance of matching a large number of pairs that share *g*.

**Fig. 3.**
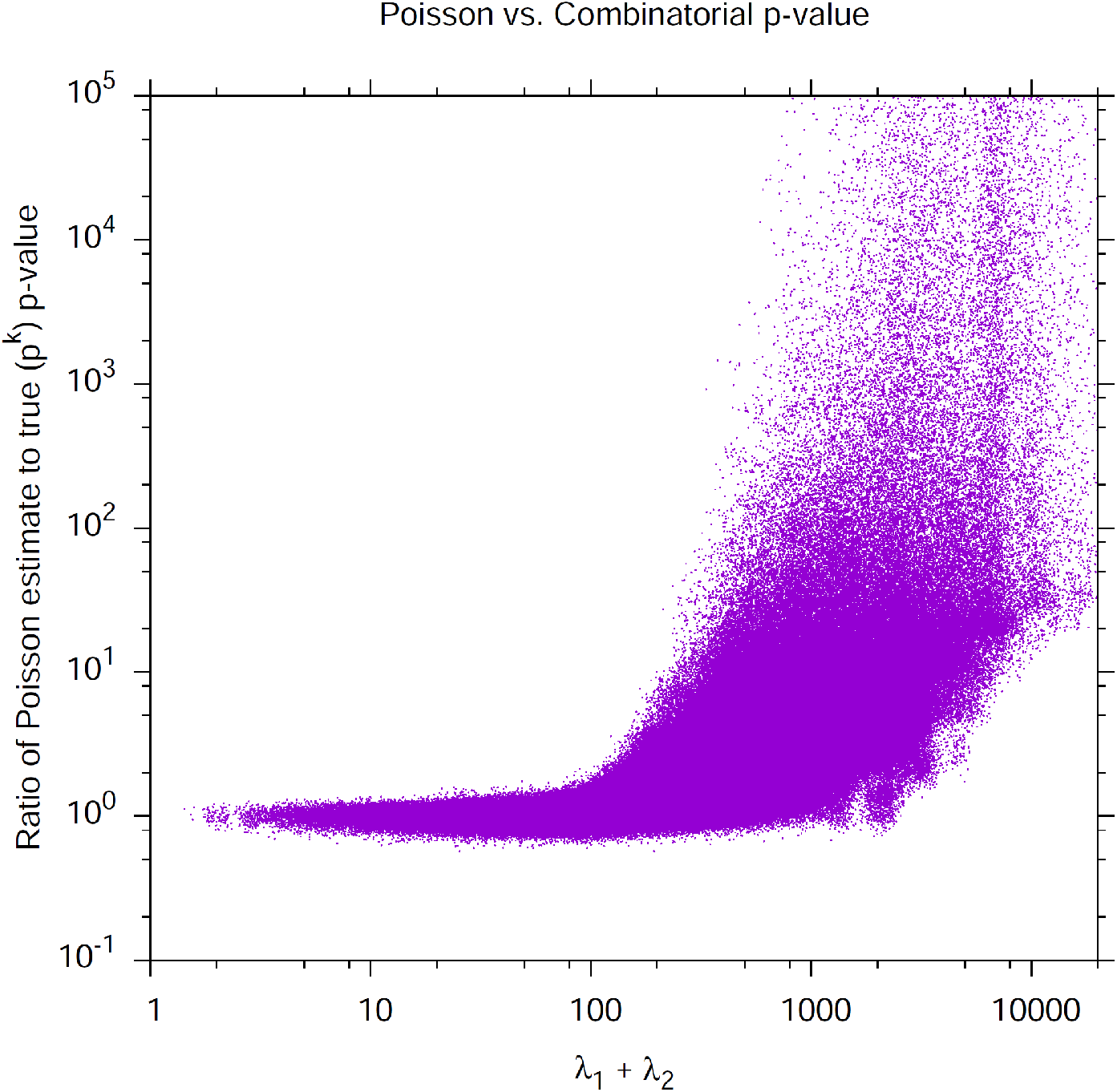
Ratio of the Poisson-based *p*-value of Eq. (10) to the exact *p*-value of Eq. (6), across 562,000 random samples from *good* alignments between networks of BioGRID 3.4.164 [3]. As we can see, the *p*-value returned by the Poisson model gets progressively worse (underestimating statistical significance) as the λ values grow. Note that some points had ratios as high as 10^70^, though we truncated the vertical axis at 10^5^. Each point has been perturbed in a random direction by a distance distributed as *N*(0,10^-1^) in log space, otherwise thousands of points land at the same integer co-ordinates, making it impossible to visualize the density of points across the plane.

## 4 Discussion

### 4.1 The fallacy of evaluating an alignment as an average across pairs of aligned nodes

There is a crucially important case that is often implicitly ignored in the literature by methods that evaluate GO-based significance of alignments by evaluating all aligned protein pairs, rather than evaluating all GO terms. This case is alluded to by phrases such as “consider the GO terms shared by a pair of aligned proteins…”. The problem is when there is a GO term *g* that exists in both networks (ie., 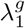 and 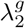 are both nonzero), but no pair of aligned proteins share it. Then the “consider…” phrase above implicitly misses the fact that *g could* have been shared by some aligned protein pairs, but was not. Unless taken care of explicitly, the alignment evaluation fails to penalize the alignment for failing to provide any matches for GO term *g*. In contrast, our method is correctly penalized for such cases: any GO term *g* for which *k* = 0 but both λ_1_ and λ_2_ are nonzero receives the appropriate penalty of a *p*-value with little statistical significance. Unfortunately, since many existing publications ignore this case, many published *p*-values claim far more statistical significance than actually exists.

A second major problem with existing *ad hoc* measures is that they do not scale even remotely monotonically with statistical significance. Take the Jaccard similarity, which is the most popular according to Table 2, though it has variously been called *GO Correctness* or *Consistency* (GOC), as well as *Functional Correctness/Consistency* (FC). Formally, given node *u* ∈ *V*_1_ aligned to *v* ∈ *V*_2_, let *S_u_, S_v_* be the set of GO terms annotating *u, v*, respectively. Then the Jaccard/GOC/FC between *u* and *v* is defined as

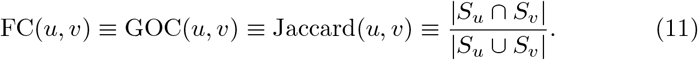

**Table 2.** Sample of published network alignment algorithm names, with their citation, year, and the method(s) they used to evaluate functional similarity. The rows are sorted by publication year; the columns are sorted by popularity of evaluation measure. **Header Legend:** Jac=Jaccard Similarity (called “GOC” and “FC” by some authors); Com=number of “common” GO terms in the cluster; MNE=Mean Normalized Entropy; Res=Resnik[4,5]; Sch=Schlicker’s method[6]; Enr=Enrichment of GO terms in a cluster compared to average cluster; *m*-sim=similarity using only GO terms with frequency (λ in our notation) less than *m*.

| Eval<br>Algo | year | Jac | Com | MNE | Res | Sch | Enr | m-sim |
| --- | --- | --- | --- | --- | --- | --- | --- | --- |
| GRAAL [9] | 2010 | . | ✓ | . | . | . | . | . |
| H-GRAAL [10] | 2010 | . | ✓ | . | . | . | . | . |
| MIGRAAL [11] | 2011 | . | ✓ | . | . | . | . | . |
| GHOST [12] | 2012 | . | . | . | ✓ | . | . | . |
| NETAL [13] | 2013 | . | ✓ | . | . | ✓ | . | . |
| SPINAL [14] | 2013 | ✓ | . | . | . | . | . | . |
| PIswap [15] | 2013 | ✓ | . | . | . | . | . | . |
| BEAMS [16] | 2014 | ✓ | . | . | . | . | . | . |
| NetCoffee [17] | 2014 | . | . | . | . | ✓ | . | . |
| MAGNA [18] | 2014 | . | ✓ | ✓ | . | . | . | . |
| HubAlign [19] | 2014 | ✓ | . | . | ✓ | ✓ | . | . |
| SiPAN [20] | 2015 | ✓ | . | . | . | . | . | . |
| FUSE [21] | 2015 | ✓ | . | ✓ | . | . | . | . |
| MeAlign [22] | 2015 | ✓ | . | . | . | . | . | . |
| OptNetAlign [23] | 2015 | ✓ | ✓ | . | . | . | . | . |
| LGRAAL [24] | 2015 | . | . | . | ✓ | . | . | . |
| WAVE [25] | 2015 | . | . | ✓ | . | . | ✓ | . |
| HGA [26] | 2016 | . | ✓ | ✓ | . | . | . | . |
| DirectedGr [27] | 2016 | . | . | . | . | . | ✓ | . |
| ModuleAlign [28] | 2016 | . | . | . | . | ✓ | . | . |
| ConvexAlign [29] | 2016 | ✓ | . | ✓ | . | ✓ | . | . |
| PROPER [30] | 2016 | ✓ | . | . | . | . | . | . |
| GMalign [31] | 2017 | ✓ | . | . | ✓ | . | . | . |
| INDEX [32] | 2017 | ✓ | ✓ | . | ✓ | . | . | . |
| Ualign [33] | 2017 | . | . | . | . | . | ✓ | . |
| SANA [34] | 2017 | . | . | . | ✓ | . | . | . |
| GLalign [35] | 2018 | . | . | . | ✓ | . | . | . |
| PrimAlign [36] | 2018 | ✓ | . | . | . | . | . | . |
| IBNAL [37] | 2018 | ✓ | . | . | . | . | . | . |
| MAPPIN [38] | 2018 | ✓ | . | ✓ | . | ✓ | . | . |
| multiMagna [39] | 2018 | . | ✓ | ✓ | . | . | . | . |
| MUNK [8] | 2019 | ✓ | . | . | . | . | . | ✓ |

Given this similarity across all aligned pairs of proteins, the entire alignment is given an FC value equal to the mean across all aligned pairs.

It is easy to construct a scenario to demonstrate how badly the Jaccard/GOC/FC measure can lead one astray. Consider the following simple system: network *G* has *n* = 1000 nodes. Each node is annotated with one, and only one, GO term. The first 900 nodes are annotated with GO term *g*_0_—ie., λ^*g*_0_^ = 900. We will refer to these as the “common” nodes. The remaining 100 nodes are each individually annotated with their own unique GO term, with names {*g*_1_, *g*_2_,…, *g*_99_, *g*_100_}; thus, λ^*g_i_*^ = 1 for all *i* = 1,…, 100. We will refer to these as the “specific” nodes. For simplicity, we will align *G* to itself, and assume that all 101 of the GO terms are *independent*, so that the *p*-value of the entire alignment is the product of the *p*-values across 101 GO terms.^1^ Then, every pair of aligned nodes constitutes a *cluster*, and the only possible per-cluster FC scores are 0 and 1, so that the mean alignment-wide FC score is simply the fraction of node pairs that are *matched*, using the formal definition of “match” from Section §2.3.

In a random alignment of G to itself, each common node has a 90% chance of being aligned with another common node, so that the expected number of matched common nodes is 810; evaluating Eq. (6) we find that *p*_810_(1000,1000, 900, 900) = 0.139—not statistically significant, as expected. On the other hand, each specific node has only a 0.1% chance of being aligned with its one and only match, so that in a random alignment we expect *none* (or very few) of the specific nodes to match. For this example, assume we do a bit better on the common nodes and match 820 of them, but match none of the specific nodes. The *p*-value of matching 820 common nodes is 0.0007. The 100 unmatched specific nodes each have *p*_0_(1000, 1000, 1, 1) = 0.999, and 0.999^100^ = 0.90. All told, this alignment has FC score of 0.82—making it look very good—and a *p*-value of about 0.0006.

Now consider a second alignment with the same FC score: we will correctly match just 10 of the specific nodes, and assume the other 90 are aligned with common nodes. This leaves precisely 810 common nodes to match each other, so the FC score is (810+10)/1000 = 0.82, as above. The *p*-value of 810 matched common nodes is above—0.139. However, each specific node has probability only 10^-3^ of matching in a random alignment, so the *p*-value of matching 10 of them is 10^-30^.

Thus, both alignments have a mean FC of 0.82, yet—to the nearest order-of-magnitude—the first has a *p*-value of only ≈ 10^-3^, while the second has a *p*-value of ≈ 10^-31^. From a statistical significance standpoint, the second one is—quite literally—*astronomically* more significant.

The takeaway message is that any method that evaluates functional significance across pairs of aligned proteins, rather than across GO terms, is likely to lead to very misleading conclusions.

### 4.2 The problem with ignoring GO terms close to the root of the hierarchy

A common practice [2] involves arbitrarily ignoring GO terms in the top few levels of the GO hierarchy on the assumption that, when a GO term annotates so many proteins, a protein pair that matches it has little value. A known problem with this suggestion [2] is the definition of “top few levels”: even GO terms at the same level but different regions of the GO hierarchy can have vastly different values of λ, so that it is difficult to choose which GO terms to ignore.

From the network alignment perspective, ignoring these common GO terms has the opposite problem to that of §4.1 in that, rather than failing to *penalize* a bad alignment, this procedure fails to adequately *reward* alignments that are “good” in the following sense. Assume a GO term *g* annotates 10% of proteins in each network. This can be a substantial number of proteins (eg., over 1700 in human and almost 700 in mouse), and such a GO term is likely to be high in the hierarchy. However, if a network alignment matches a substantially larger fraction of this plethora of pairs than is expected at random, it is a sign that *large regions* of functional similarity are being correctly aligned to each other, even if individual proteins are not. In other words, perhaps similar pathways are being correctly mapped to each other even if the individual proteins in the pathway are incorrectly mapped. A network alignment that succesfully matches such large regions should be rewarded for doing so, but if “common” GO terms are disregarded, this won’t happen.

### 4.3 Practical details regarding treatment of GO terms

Our analysis is easily adapted to evaluate network alignments based on any subset of GO terms. For example, one may wish to separately evaluate the three GO hierarchies of *Biological Process* (BP), *molecular Function* (MF), and *Cellular Component* (CC). Similarly, one should evaluate an alignment without the use sequence-based GO terms if sequence played any part in constructing the alignment.

Once we compute a rigorous *p*-value for each GO term *g* that appears in both networks, computing a GO-based *p*-value of the entire alignment requires combining the multitude of “per-GO-term” *p*-values into a “holistic” GO-based *p*-value for the entire alignment. Doing so rigorously is nontrivial, since GO terms are arranged in a hierarchy and are not independent of each other. While we provide here the first rigorous measure of the *p*-value of an alignment with respect to an individual GO term, to our knowledge nobody has yet worked out how to rigorously account for the inter-dependence of GO terms. We leave for future work the job of providing such a holistic *p*-value.

## Appendix: brief survey of existing GO-based measure of network alignments

Table 2 presents a list of alignment papers and the measures they use to evaluate functional similarity. Without exception, all of these methods evaluate each pair of aligned nodes individually, and then take some sort of average across pairs. (Some methods are not 1-to-1 and so the “pair” of aligned nodes we discuss must be generalized to a *cluster* of aligned nodes, but this generalization does not negate our point.) We are aware of no existing methods that consider the alignment from the perspective of one GO term’s performance globally across all clusters. Thus, all of these methods suffer the major drawbacks described in Section §4.1.

Below is a brief description of the methods.

– **Jaccard / FC / GOC:** A common pairwise measure is the Jaccard similarity of Eq. (11), often called GOC (for GO “consistency” or “correctness”) or FC (functional consistency/correctness).
– **Common GO terms:** In a PGNA, choose an integer threshold *h* (usually 1–5), and count how many aligned pairs have at least h GO terms in common. No effort is made to account for the annotation frequency (λ in our notation) of any GO term.
– **Entropy:** Given a cluster of proteins *S* in which *d* GO terms {*g*_1_,…, *g_d_*} appear at least once across all the proteins in *S*, the entropy is defined as 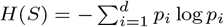, where *p_i_* is the fraction of all proteins in *S* that are annotated with GO term *g_i_*. Entropy is always non-negative and lower values are better. The *normalized entropy* is *N*(*S*) = *H*(*S*)/*m*, where *m* is the number of unique GO terms in S. Alignments can then be scored using *Mean Entropy* (ME) or *Mean Normalized Entropy* (MNE), which is just the appropriate mean across all clusters *S*.
– **Resnik**: Based on Resnik’s measure of semantic similarity [4,5], it was originally designed only to evaluate the similarity between two GO terms by finding their *most informative common ancestor* in the GO hierarchy, and using an information-theory argument to compute their common information. Later it was extended to measure similarity between gene products, such as proteins, by taking some sort of mean or maximum between the GO terms of two proteins (see, eg., [6,7,2].
– **Schlicker’s method [6]**: an extension of Resnik’s measure tailored more closely to genes and gene products.
– **Enrichment**: has been defined in various ways but usually measures whether the shared GO terms between a pair of aligned proteins (or more generally in a cluster) is “enriched” beyond what’s expected at random.
– *m***-sim**: This measure is only used by MUNK [8], which is not technically a network alignment algorithm, though it is designed to find functionally similar genes or proteins between species. It is also the only method from Table 2 that takes into account the frequency of annotation of a GO term *g* (λ^*g*^ in our notation), by using only GO terms with λ below some threshold *m*.

## Compliance with Ethical Standards

This work was unfunded, and the author declares no competing interests.

## Footnotes

1 The assumption of independence is not entirely unfounded; for example we could choose go to be the *Cellular Component* (CC) GO term `GO:0005634`, which describes the location “nucleus”, and choose the remainder of GO terms to be *molecular functions* (MF) that tend to occur only outside the nucleus. In fact, in the Sept. 2018 release of the GO term database there are over 700 MF GO terms with the following properties: (a) they annotate exactly one protein (ie., each of over 700 GO terms *g* has λ^*g*^ = 1), and (b) for each such GO term, the one protein it annotates is *not* annotated with `GO:0005634`. The fact that over 700 such GO terms exist make our independence assumption plausible—at least in this artificial scenario.

